# Deep sequencing of pre-translational mRNPs reveals hidden flux through evolutionarily conserved AS-NMD pathways

**DOI:** 10.1101/847004

**Authors:** Carrie Kovalak, Scott Donovan, Alicia A. Bicknell, Mihir Metkar, Melissa J. Moore

## Abstract

**Background:** Alternative splicing (AS), which generates multiple mRNA isoforms from single genes, is crucial for the regulation of eukaryotic gene expression. The flux through competing AS pathways cannot be determined by traditional RNA-Seq, however, because different mRNA isoforms can have widely differing decay rates. Indeed, some mRNA isoforms with extremely short half-lives, such as those subject to translation-dependent nonsense-mediated decay (AS-NMD), may be completely overlooked in even the most extensive RNA-Seq analyses.

**Results:** RNA immunoprecipitation in tandem (RIPiT) of exon junction complex (EJC) components allows for purification of post-splicing mRNA-protein particles (mRNPs) not yet subject to translation (pre-translational mRNPs) and, therefore, translation-dependent mRNA decay. Here we compared EJC RIPiT-Seq to whole cell RNA-Seq data from HEK293 cells. Consistent with expectation, the flux through known AS-NMD pathways is substantially higher than that captured by RNA-Seq. Our EJC RIPiT-Seq also definitively demonstrates that the splicing machinery itself has no ability to detect reading frame. We identified thousands of previously unannotated splicing events; while many can be attributed to “splicing noise”, others are evolutionarily-conserved events that produce new AS-NMD isoforms likely involved in maintenance of protein homeostasis. Several of these occur in genes whose overexpression has been linked to poor cancer prognosis.

**Conclusions:** Deep sequencing of RNAs in post-splicing, pre-translational mRNPs provides a means to identify and quantify splicing events without the confounding influence of differential mRNA decay. For many known AS-NMD targets, the NMD-linked AS pathway predominates. EJC RIPiT-Seq also enabled identification of numerous conserved but previously unannotated AS-NMD events.

## Background

A central mechanism underlying metazoan gene expression is alternative pre-mRNA processing, which regulates the repertoire of mRNA isoforms expressed in various tissues and under different cellular conditions. Extensive deep sequencing of RNA (RNA-Seq) has revealed that ∼95% of human protein-coding genes are subject to alternative splicing (AS) [1, 2], with current estimates suggesting ∼82,000 different protein-coding mRNA isoforms generated from ∼20,000 protein coding genes [3]. Thus, production of alternative mRNA isoforms massively expands the protein repertoire that can be expressed from a much smaller number of genes [4, 5]. But cells also need to control how much of each protein is made. Although transcriptional control is often considered the predominant mechanism for modulating protein abundance, emerging evidence indicates that post-transcriptional regulatory mechanisms are crucial as well.

Not all mRNA variants are protein-coding. Nearly 15,000 human mRNAs in the Ensembl database (release 93) are annotated as nonsense-mediated decay (NMD) targets [3]. NMD is a translation-dependent pathway that both eliminates aberant mRNAs with malformed coding regions (i.e., those containing premature termination codons due to mutation or missplicing) and serves as a key mechanism for maintenance of protein homeostasis [6]. This protein homeostasis function is mediated by AS linked to NMD (AS-NMD), wherein the flux through alternate splicing pathways that result in protein-coding and NMD isoforms is subject to tight control [7]. These NMD isoforms harbor a premature termination codon either due to frameshifting or inclusion of a poison cassette exon. Because NMD isoforms are rapidly eliminated after the first or “pioneer” round of translation, only protein-coding isoforms result in appreciable protein production (**Figure 1A, bottom**). Thus increasing or decreasing flux through the NMD splicing pathway decreases or increases protein production, respectively. Although AS-NMD was originally described as a mechanism by which RNA binding proteins (e.g., SR and hnRNP proteins) could autoregulate their own synthesis, recent work indicates that AS-NMD is much more pervasive, tuning abundance of many other proteins such as those involved in chromatin modification and cellular differentiation [8].

**Figure 1.**
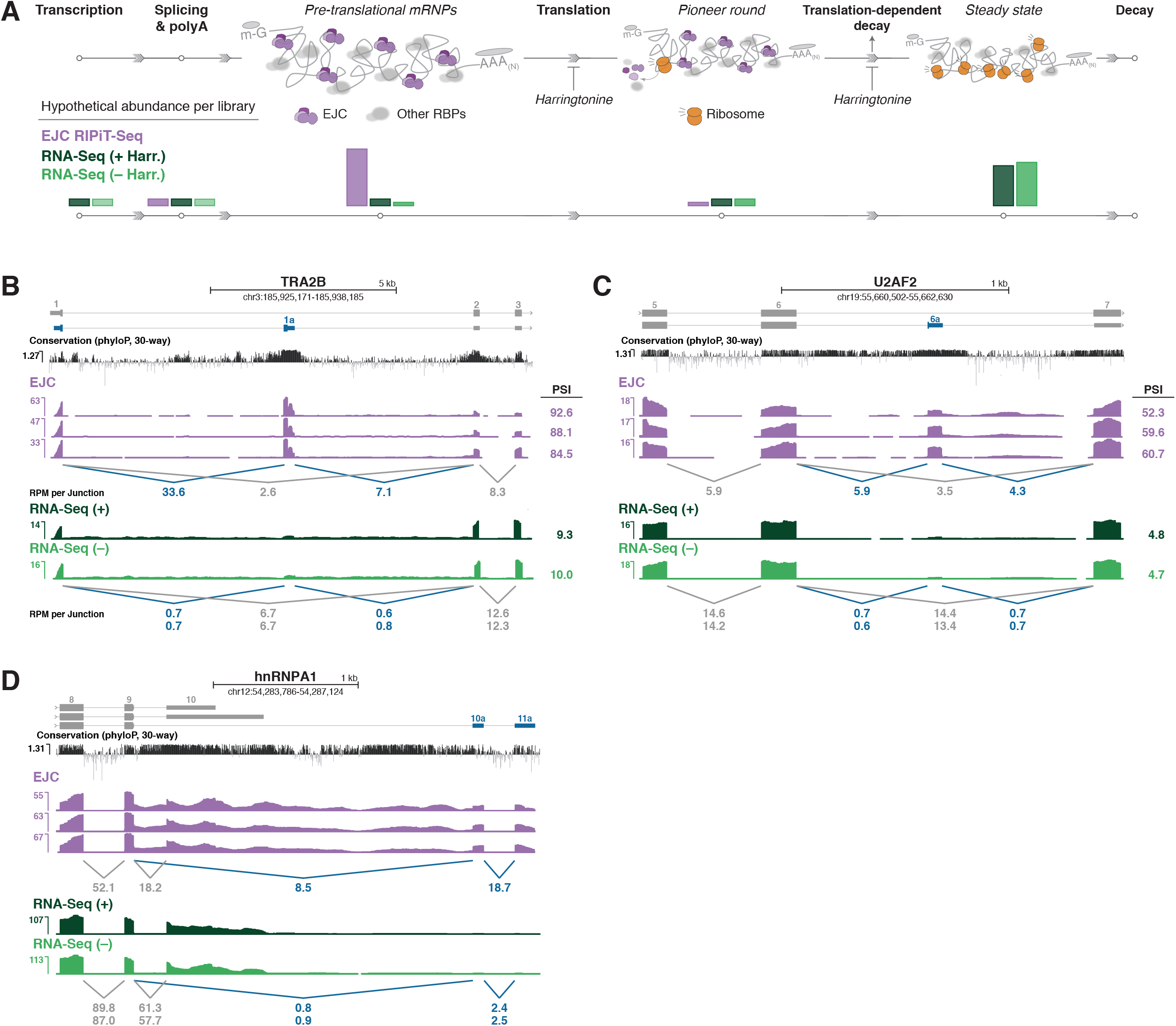
(A) (Top) mRNA metabolism from transcription to degradation. EJCs (purple) deposited upstream of exon junctions and other RNA-binding proteins (RBPs; grey) are cleared by ribosomes (orange) during the pioneer round of translation. While Protein-coding isoforms are subject to multiple rounds of translation prior to decay, NMD isoforms are rapidly eliminated. Steps affected by harringtonine treatment are indicated. (Bottom) Hypothetical abundance throughout the mRNA lifecycle in the libraries analyzed in this paper: EJC-bound RIPiT-Seq (purple) and RNA-Seq libraries treated with (+; dark green) or without (–; light green) harringtonine. (B-D) Genome browser tracks of library coverage across individual genes (grey: protein-coding isoform(s); blue: NMD isoform) containing poison cassette exons (B, TRA2B and C, U2AF2) or 3′ UTR introns (D, hnRNPA1). Shown are all three EJC RIPiT-Seq replicates and replicate 1 for both (+) and (–) harringtonine RNA-Seq libraries. Conservation tracks show phyloP basewise scores derived from Multiz alignment of 30 vertebrate species. Numbers below tracks indicate mean reads per million (RPM) spanning each exon junction. Numbers on the right in B and C are percent spliced in (PSI) values for poison exon inclusion events; PSI values for RNA-Seq libraries are replicate means. See Methods for PSI formula.

The true extent to which AS-NMD contributes to protein homeostasis can only be appreciated by determining flux through the splicing pathways that alternately produce protein-coding and NMD isoforms. Transcriptome-wide assessment of mRNA isoform abundance generally relies on RNA-Seq, which provides a static snapshot of the species present in the sample at the time of collection. Because NMD isoforms are so rapidly decayed, they are generally underrepresented in RNA-Seq datasets. Thus a single RNA-Seq snapshot is generally uninformative as to synthetic flux through protein-coding and NMD splicing pathways.

An alternate means to assess protein-coding and NMD pathway flux is to capture newly synthesized mRNAs after splicing completion but prior to translation. Late in the splicing cycle, the exon junction complex (EJC) is deposited upstream of at least 80% of exon-exon junctions (canonical; cEJCs) and at multiple other sites throughout the length of spliced exons (noncanonical; ncEJCs) [9, 10]. Upon nucleocytoplasmic export, the pioneer round of translation removes EJCs within the 5′ UTR and CDS regions, with EJCs remaining downstream of stop codons being key mediators of NMD [11]. Pre-translational mRNPs can be selectively isolated by tandem immunoprecipitation of epitope-tagged and untagged EJC components, a technique known as RNA:protein immunoprecipitation in tandem (RIPiT) [12]. Deep sequencing library preparation from RIPiT samples (RIPiT-Seq) has previously enabled us to map the positions of canonical and noncanonical EJCs on spliced transcripts [9] and to investigate the RNA packing principles within pre-translational mRNPs [13].

Here, we compare libraries from pre-translational mRNPs (EJC RIPiT) isolated from subconfluent HEK293 cells to matched RNA-Seq libraries (**Figure 1A**). As expected, EJC RIPiT-Seq libraries are enriched for transcript isoforms destined for translation-dependent decay. By providing a window into the repertoire of transcripts generated by splicing but prior to translation-dependent decay, EJC RIPiT-Seq libraries provide a more accurate record of the flux through various alternative processing pathways than does standard RNA-Seq. Importantly, EJC RIPiT-Seq libraries enabled us to identify numerous new evolutionarily-conserved poison cassette exons that had previously eluded annotation.

## Results

### EJC and RNA-Seq libraries

In our recent study investigating the organizing principles of spliced RNPs [13], we generated three biological replicates from subconfluent HEK293 cells of EJC-bound RNAs partially digested with RNase T1 during RNP purification (**Figure 1A**). Paired-end deep sequencing of these EJC RIPiT-Seq libraries resulted in 19-25 million mate pairs each (**Supplemental Table 1**). To enable comparison to RNA-Seq for the current study, we created and sequenced rRNA-depleted whole cell RNA-Seq libraries (84-93 million mate pairs each) wherein the captured fragments were of similar length (220-500 nts) to our previously published EJC RIPiT-Seq libraries. The new RNA-Seq libraries were generated from cultures (three biological replicates each) that were (+) or were not (–) subjected to a one hour pre-treatment with harringtonine. Harringtonine, a translation initiation inhibitor, is used in the EJC RIPiT-Seq protocol to enrich for pre-translational mRNPs [9, 12, 60].

For all libraries, raw reads were aligned to the Genome Reference Consortium Human Build 38 (GRCh38.p12) [3] using STAR (v2.5.3a) [14] after first filtering out those mapping to repeat RNAs [15]. To minimize any effects due to misalignment in ensuing analyses, mismatches were limited to three per read, with gaps caused by deletions or insertions being strongly penalized. These strict mapping parameters resulted in 6-10 million and 60-76 million aligned pairs for the EJC RIPiT-Seq and RNA-Seq libraries, respectively (**Supplemental Table 1**). For quantification, we limited all analyses to unique reads with high mapping quality (MAPQ ≥ 5). For all libraries, we used Kallisto (v0.44.0) to derive expression values for the ∼200,000 annotated transcripts in GRCh38.p12 [3]. Examination of per-transcript abundance revealed high concordance (≥ 0.93 to 0.99) among biological replicates (**Supplemental Figure 1A**). Therefore, all subsequent quantitative analyses utilized merged biological replicate data.

### EJC libraries are enriched for spliced transcripts and translation-dependent decay targets

To assess the relative abundance of NMD targets in EJC and RNA-Seq libraries, we first examined read coverage on known AS-NMD genes. The SR proteins TRA2B and U2AF2 negatively regulate their own expression by promoting inclusion of a highly-conserved poison cassette exon containing a premature termination codon (**Figure 1B, C**). Although these poison exons were detectable in all library types, they were substantially more abundant in the EJC libraries. Whereas the RNA-Seq libraries returned low poison exon inclusion values (percent spliced in; PSI; 9.3-10.0% and 4.7-4.8%, respectively), the EJC RIPiT-Seq libraries indicate much higher inclusion rates (88.4% and 57.5%, respectively). Thus, for both TRA2B and U2AF2, the predominant splicing pathway in HEK293 cells under standard growth conditions is poison exon inclusion. Similar trends were observed for other known AS-NMD targets (**Supplemental Figure 1B-D**), including hnRNPA1 where the AS-NMD isoform results from 3′ UTR splicing as a consequence of alternative polyadenylation (**Figure 1D**). Importantly, the (–) and (+) harringtonine RNA-Seq libraries exhibited nearly identical AS-NMD isoform abundances. Thus a 60 minute inhibition of translation was insufficient to substantially change AS-NMD isoform abundance in whole cell RNA-Seq libraries from subconfluent HEK293 cells. In contrast, the substantial differences between the EJC RIPiT-Seq and RNA-Seq quantitations for these previously documented AS-NMD isoforms clearly illustrate the advantage provided by EJC RIPiT-Seq for more accurately assessing flux through alternative processing pathway resulting in mRNA isoforms with widely different decay rates.

In GRCh38.p12, every transcript isoform is given a specific annotation [16]; relevant annotations in protein-coding genes are “protein-coding”, “NMD”, “NSD”, “retained intron”, and “processed transcript”, with the latter being a catch-all for transcripts not clearly attributable to any other category. NSD (non-stop decay) is another translation-dependent mRNA degradation pathway that eliminates transcripts having no in-frame stop codon [17]. Retained intron transcripts are generally subject to translation-dependent decay driven by in-frame stop codons in the intronic regions. For transcripts detectable in our libraries [TPM >0 in all replicates of a particular library type: EJC and (+) or (–) RNA-Seq], the number of exon junctions (i.e., positions at which introns were removed) ranged from 0 to >100 per protein-coding isoform and 1 to 69 per NMD isoform (**Figure 2A**). As expected, protein-coding isoforms having no exon junctions were less abundant in the EJC libraries than in RNA-Seq libraries (**Figure 2B, top**). In contrast, spliced protein-coding isoforms containing 5 or more exon junctions were enriched in EJC libraries, with the degree of enrichment increasing with exon junction number. For each exon junction number bin (i.e., 1-4, 5-10 and 10+), NMD isoforms were even more enriched in EJC libraries than were protein-coding isoforms (**Figure 2B, bottom**). EJC library enrichment was also readily discernible for NMD, NSD, retained intron, and processed transcript isoforms (**Figure 2C** and **Supplemental Figure 2A**), with median enrichments falling between 1.8 and 2.6-fold (**Supplemental Figure 2B**). Because the high degree of overlap between alternate transcript isoforms from individual genes confounds individual isoform abundance quantification by algorithms such as Kallisto [18, 19], we performed an additional analysis examining only those exon junctions (or intron-exon boundaries for retained introns) not shared between multiple GRCh38.p12 transcripts. This revealed an even greater enrichment of NMD, NSD, retained intron, and processed transcript isoforms in EJC than RNA-Seq libraries (**Figure 2D**), with median fold enrichments ranging from 2.5 to 2.9-fold **(Figure 2E**; note that the number of NSD transcripts with unique exon junctions was insufficient to provide statistical significance). Thus EJC RIPiT-Seq libraries are highly enriched for spliced transcripts subject to rapid clearance by translation-dependent decay.

**Figure 2.**
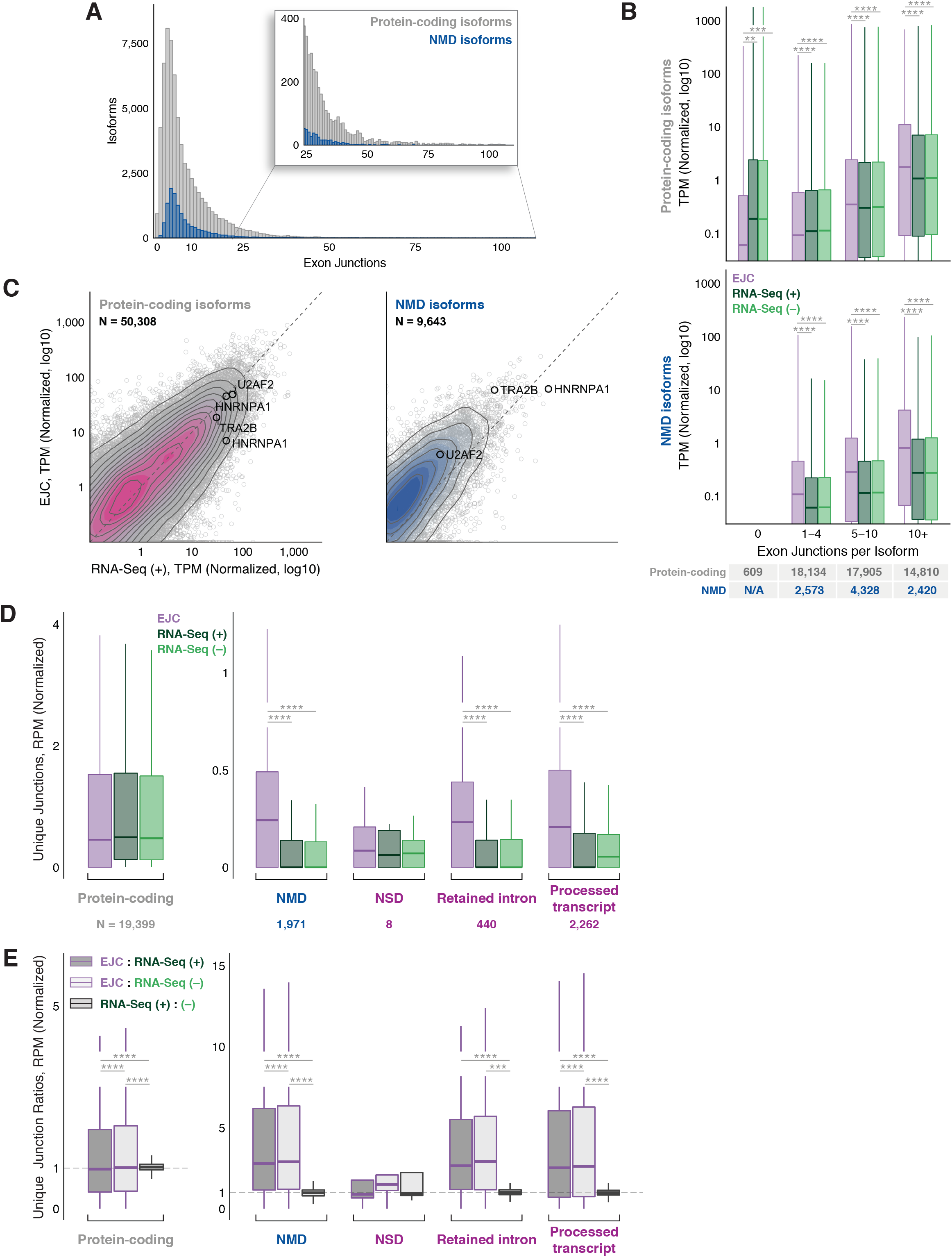
(A) Distribution of the number of exon junctions in all annotated protein-coding (grey) or NMD (blue) transcripts. (B) Distribution of protein-coding (top) and NMD (bottom) transcripts per million (TPM) in each library type (colors as in Figure 1A), binned based on indicated number of exon junctions per transcript. Numbers in bottom table: Total number of expressed isoforms per exon junction number bin. (C) Scatterplots comparing TPMs between EJC RIPiT-Seq and RNA-Seq (+) libraries for protein-coding (left) and NMD (right) isoforms. Transcripts from Figure 1 are noted. N: Number of detected transcripts (out of all annotated transcripts of that type). Dashed black line: x=y. (D) Distribution of read counts at unique junctions, binned based on transcript biotypes [3]. Numbers below: Total number of unique junctions detected in our libraries per transcript biotype. (E) Ratios of read counts at unique junctions between two indicated library types, binned based on transcript biotype [3]. For B, D and E: Results of one-way ANOVA and Tukey’s *post hoc* significance tests comparing EJC RIPiT-Seq to RNA-Seq libraries are indicated as *P<0.05, **P<0.01, ***P<0.005, ****P<0.0001.

### EJC libraries capture previously unannotated exon junctions

We next wondered whether EJC libraries might contain transcript isoforms that had heretofore escaped annotation due to their low abundance in RNA-Seq. Such isoforms should contain previously unannotated exon junctions. To identify all annotated exon junctions, we integrated the RefSeq (hg38) [20], Ensembl (GRCh38.p12) [3], GENCODE (v29) [21] and Comprehensive Human Expressed SequenceS (CHESS) transcriptome annotations to create a comprehensive reference file containing 575,837 known introns (**Supplemental Table 2**). While Ensembl and GENCODE are largely identical [62], our analysis revealed 240 junctions in Ensembl GRCh38.p12 that were not in GENCODE v29, and 4,554 junctions in GENCODE v29 that were not in Ensembl GRCh38.p12 (**Supplemental Table 2**). CHESS is derived from 9,795 RNA-Seq samples from diverse cell types in the GTEx collection, so represents the most complete compendium of human transcripts reported to date [22]. Nonetheless, while CHESS found 118,043 new exon junctions not previously annotated in RefSeq, Ensembl or GENCODE, 106,223 other junctions present in RefSeq, Ensembl and/or GENCODE were not returned by the CHESS pipeline (**Figure 3A**). This lack of concordance shows that even the most comprehensive RNA-Seq data analyses are unlikely to annotate all bona fide splicing events.

**Figure 3.**
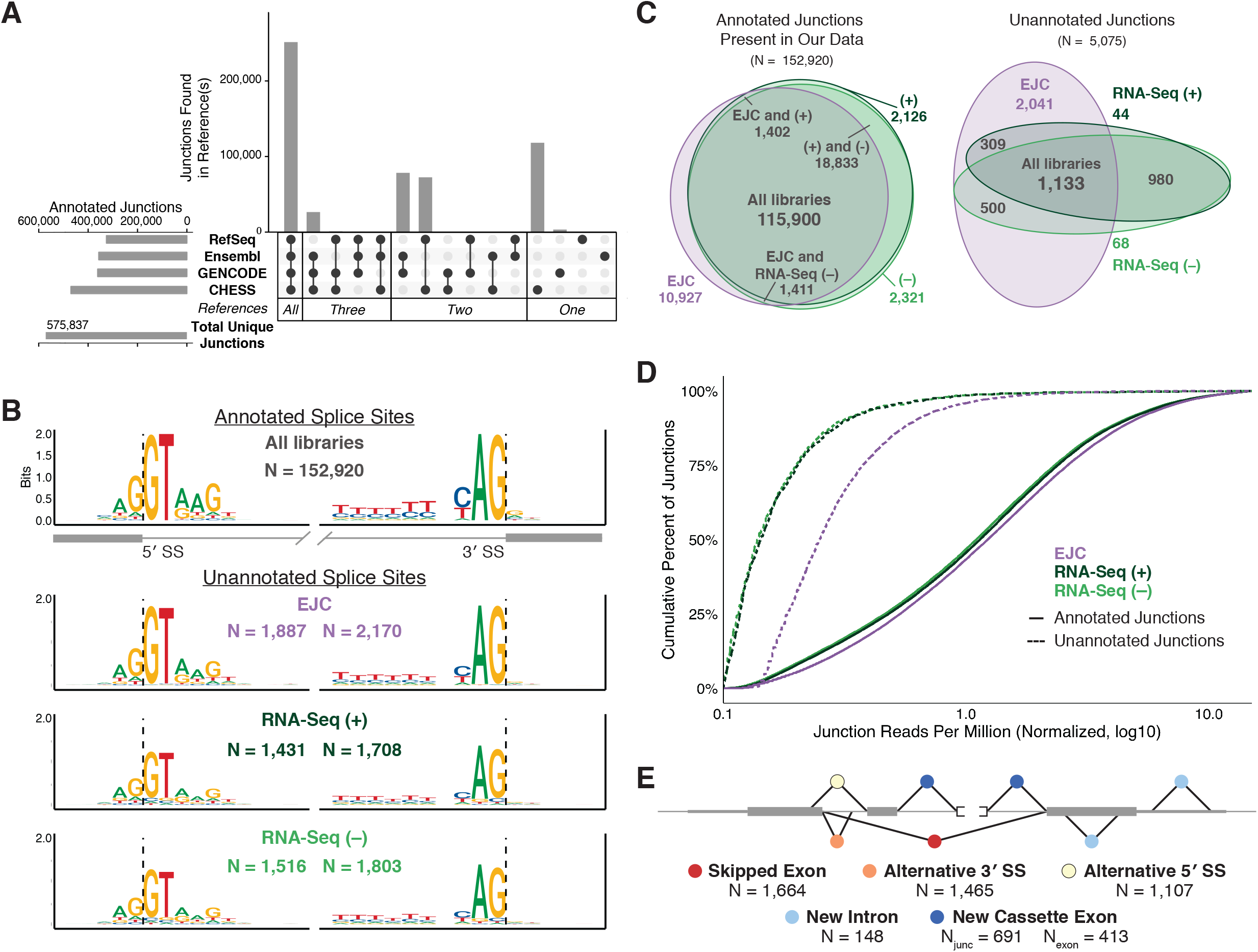
(A) Comparison of annotated exon junctions among the transcriptomes sourced from RefSeq (hg38), Ensembl (GRCh38.p12), GENCODE (v29), and CHESS (v2.1). Horizontal bars: total junctions in each reference set; vertical bars: intersections of indicated reference sets. Bar graphs created with UpSetR [56]. (B) Sequence motifs for 5′ (left) and 3′ (right) splice sites used in annotated junctions observed in at least one analyzed library type (top) and for previously unannotated splice sites in indicated library type (bottom). Sequence logos were generated in R using ggseqlogo [57]; letter height signifies the relative abundance of that nucleotide at each position. N: Number of splice sites contributing to each logo. Note that the number of unannotated junctions (6,363) is greater than the total number of unannotated splice sites because many unannotated junctions combine an annotated and unannotated splice site (i.e., alternative 5′ or 3′ splice sites). (C) Venn diagram of annotated and previously unannotated junctions (numbers indicated) shared between library types. Venn diagrams made with eulerr [58]. (D) Cumulative histogram of exon junction reads (RPM) at annotated (solid line) and previously unannotated (dashed line) junctions in each library type (colors as in Figure 1A). (E) Schematic of unannotated splicing events separated by event type: Skipped exon (red); alternative 3′ (orange) or 5′ (yellow) splice site; new intron (light blue); new cassette exon (dark blue). N: number of observed events; for new cassette exons, both the number of observed unannotated junctions and number of new exons are shown.

To identify annotated and unannotated exon junctions in our EJC and RNA-Seq libraries, we considered only those reads that cross an exon junction. The position of an exon junction in an individual read can be found by examining the “N operation” in the CIGAR string, which indicates the locations and lengths of gaps inserted during alignment to genomic DNA (**Supplemental Figure 3A**). We further required that any candidate junction: (1) occur within an annotated gene; (2) have reads with ≥15 nt aligning on both sides of the junction (≥90% exact sequence match on each side); (3) be detectable in all replicates of a particular library type; and (4) have a mean read count ≥2 per library type (**Supplemental Figure 3A**). Using these criteria, we identified 152,920 junctions contained in the RefSeq/Ensembl/GENCODE/CHESS reference file (annotated junctions) and 6,363 unannotated junctions. MEME analysis of the latter revealed the 5′ and 3′ splice site consensus motifs for the major spliceosome, although at somewhat lesser strength (bits) than annotated junctions (**Figure 3B**). To limit our analysis to events most likely representing real splicing events (as opposed to mapping artifacts), we subsequently only considered the 5,075 previously unannotated junctions where the putative intron began and ended with dinucleotides GT-AG or AT-AC to include excision events mediated by both the major and minor spliceosomes (**Supplemental Table 3**).

The majority (76%) of previously-annotated exon junctions meeting our detection criteria in protein coding genes (**Supplemental Figure 3A**) were present in all three library types (**Figure 3C, left**). There was less concordance, however, with respect to unannotated junctions, with the EJC libraries having many more unannotated junctions than either (+) or (–) harringtonine RNA-Seq (**Figure 3C, right**). Consistent with this, unannotated junctions were supported by more reads per million mapped (RPM) in EJC than in either RNA-Seq library (**Figure 3D**; p=2.2E-16, Kolmogorov–Smirnov test), while read coverages over annotated junctions were remarkably similar between library types. The major class (51%) of the new junctions were new alternative 5′ or 3′ splice sites (i.e., that combined a known 3′ or 5′ splice site with a previously unannotated 5′ or 3′ splice site, respectively) (**Figure 3E**). Other categories were previously unannotated exon skipping events (33%), new cassette exons (14%) and new introns (3%).

### Relationship of new splicing events to reading frame

Previous analyses of low abundance, unannotated splicing events in RNA-Seq data have revealed a strong tendency for such events to maintain reading frame [23, 24]. To investigate whether this is due to some inherent ability of the splicing machinery to detect reading frame in the nucleus [25, 26], or simply due to translation-dependent decay of out-of-frame events, we determined the distance from each previously unannotated splice site meeting our selection criteria to the nearest annotated splice site observed in any of our three library types. In all, 250 and 522 unannotated 5′ and 3′ splice sites, respectively, occurred within 15 nts of an annotated 5′ or 3′ splice site. Comparison of unannotated-to-annotated splice site distance aggregation plots between the three library types revealed both similarities and differences (**Figure 4A**). Around annotated 5′ splice sites, all three libraries displayed similar patterns, with the greatest unannotated usage being at intron position +5, consistent with the preference for a G and a T at positions +5 and +6, respectively, in the human 5′ splice site consensus sequence (**Figure 3B**) and the prevalence of GT dinucleotides at this position in this set of 250 5′ splice sites (dotted gray line in **Figure 4A**). More notable was the pattern near 3′ splice sites, where positions +3, +4 and +5 in the downstream exon exhibited the highest unannotated usage. Strikingly, whereas the RNA-Seq libraries were strongly skewed toward position +3, all three positions were highly represented in the EJC libraries, with their usage more reflective of the number of available AG’s at these positions (dotted gray line in **Figure 4A**). Comparison of fractional abundance [unannotated read counts/(unannotated + annotated read counts)] at individual sites confirmed that whereas the EJC and RNA-Seq libraries exhibited similar utilization at position +3, utilization of positions +4 and +5 was much more prominent in the EJC than either RNA-Seq library (**Figure 4B**). These observations strongly support a model in which out-of-frame splicing events are rapidly eliminated by NMD, resulting in their underrepresentation in RNA-Seq libraries. Because downstream AG utilization in the EJC libraries so closely paralleled their availability, we conclude that (at least with regard to 3′ splice sites) the splicing machinery has no ability to read frame.

**Figure 4.**
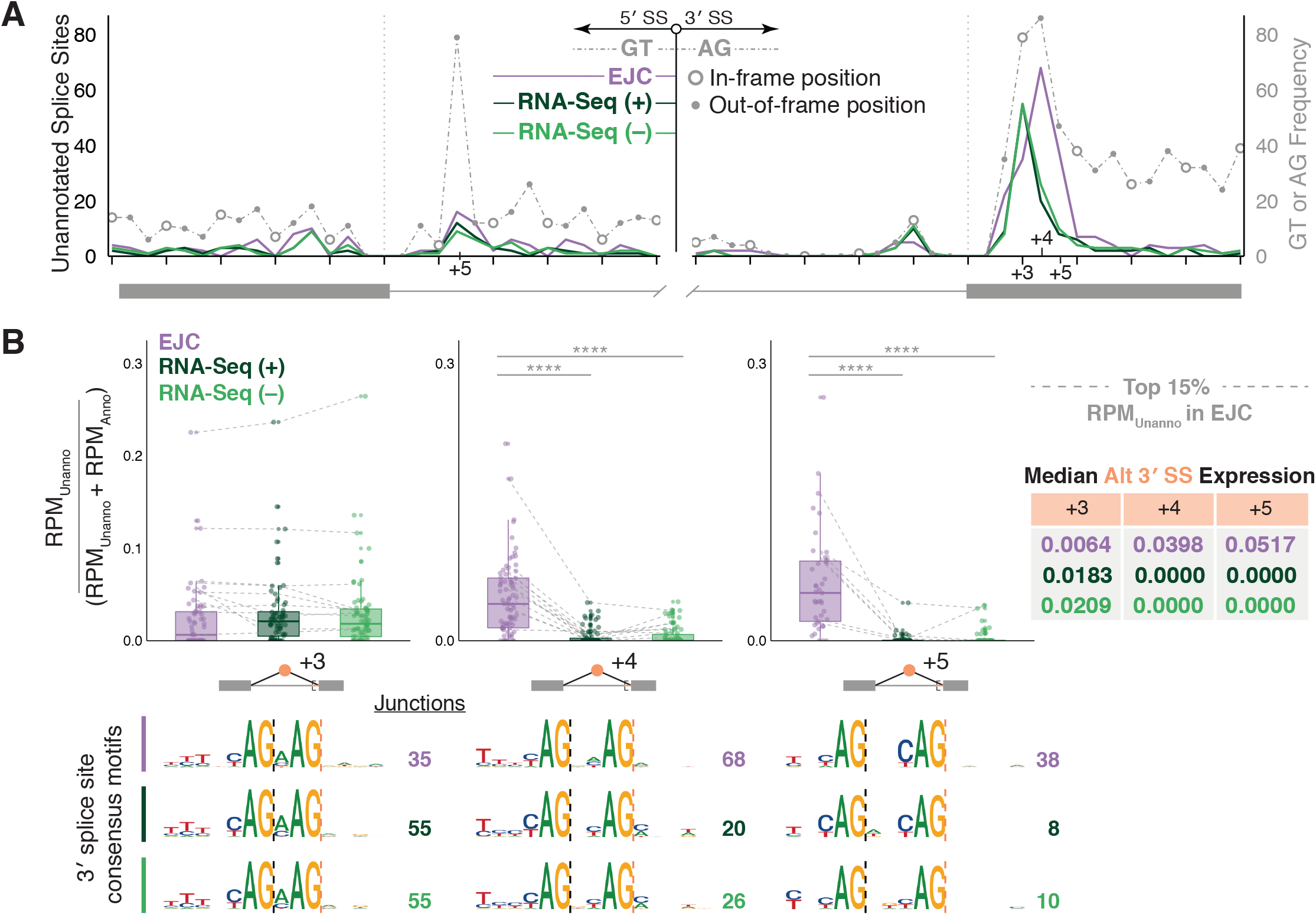
(A) Distribution of unannotated splice sites relative to the closest annotated splice site observed in analyzed libraries (solid colored lines). Grey dotted line: Frequency of available GT or AG dinucleotides surrounding the annotated 5′ (left) and 3′ (right) splice sites with open circles indicating in-frame positions and solid grey dots indicating out-of-frame positions. (B) Distribution of the ratio of unannotated alternative 3′ splice site use (RPM_Unanno_) over all events using the same 5′ splice site (RPM_Unanno_+ RPM_Anno_) in each library type. (Left) Unannotated alternative 3′ splice sites at positions +3, +4, and +5 relative to closest annotated 3′ splice site. Grey lines show how the top 15% (highest RPM_Unanno_) of unannotated junctions detected in EJC RIPiT-Seq libraries differ between library types. Results of one-way ANOVA and Tukey’s *post hoc* tests comparing EJC RIPiT-Seq to RNA-Seq libraries are indicated; ****P<0.0001. (Right) Median [RPM_Unanno_/(RPM_Unanno_+ RPM_Anno_)] values per library at the +3, +4, and +5 positions. (Bottom) Sequence motifs for unannotated 3′ splice sites used at positions +3, +4, and +5. Sequence logos were generated in R using ggseqlogo [57]; letter height signifies the relative abundance of that nucleotide at each position. N: Number of splice sites contributing to each logo. Dashed lines indicate location of annotated (black) and unannotated (orange) splice sites.

### Evolutionary conservation versus splicing noise

Regardless of reading frame, most unannotated splicing events are likely due to “splicing error” [27] or “splicing noise” [24]. Splicing noise results from spurious utilization of cryptic splice sites that are not evolutionarily conserved. To assess both evolutionary conservation and splice site strength, we calculated mean basewise phyloP 30-way vertebrate conservation [28] and MaxENT (a generally accepted measure of how well a particular splice site matches the consensus) [29] scores for both annotated and unannotated splice sites, using the same 5′ and 3′ splice site window sizes (9 and 23 nts, respectively) for both calculations (**Figure 5A**). We also calculated conservation and MaxENT scores for sequences chosen at random from inside annotated genes and containing either GT or AG at the appropriate position within the 5′ or 3′ splice site window, respectively. Plotting MaxENT versus conservation revealed markedly different distributions between annotated splice sites and random GT- and AG-containing sequences (**Figure 5B and Supplemental Figure 4**), with annotated sites being notably skewed toward higher values for both measures. In contrast, whereas unannotated splice sites were similarly distributed as annotated splice sites with regard to MaxENT, the majority exhibited conservation scores more similar to random than annotated splice sites (**Figure 5C and Supplemental Figure 4C**). For the random sequences, 95% had 5′ and 3′ splice site conservation scores below 1.04 and 0.63, respectively. Using these values as cutoffs to filter out the majority of events likely due to splicing noise (although this may be unnecessarily conservative for 3′ splice sites due to the high degree of overlap between the annotated and random conservation scores) left us with 453 (23%) and 651 (28%) evolutionarily-conserved unannotated 5′ and 3′ splice sites, respectively (**Supplemental Table 3**). The majority of these occurred within annotated protein-coding exons, so their conservation is likely driven by amino acid conservation and not as a requirement for recognition by the splicing machinery (see **Supplemental Figure 5A** for an example). Almost all of the new evolutionarily conserved introns (i.e., both the 5′ and 3′ splice sites were previously unannotated, but exhibited high conservation) also fell into this category. For the new introns, calculation of percent intron retention (PIR) in the EJC libraries revealed highly inefficient splicing (mean PIR = 84%); individual examination of those exhibiting the highest number of exon junction reads in the EJC libraries led to no findings of particular note. Thus the new introns likely constitute splicing noise due to low level spliceosome assembly on sites within exons that by happenstance resemble splice site consensus sequences. In contrast, examination of unannotated 3′ splice sites occurring within introns uncovered a conserved alternative splicing event in the HECTD4 (HECT domain E3 ubiquitin protein ligase 4) gene that adds 9 amino acids into the middle of the protein (**Supplemental Figure 5B**); this spliced isoform is currently annotated in mouse RefSeq and GENCODE, but not in humans. Other alternative 3′ splice sites in the CNOT1 and EEA1 genes generate AS-NMD isoforms (**Figure 5E, F**), the latter due to creation of a new poison cassette exon.

**Figure 5.**
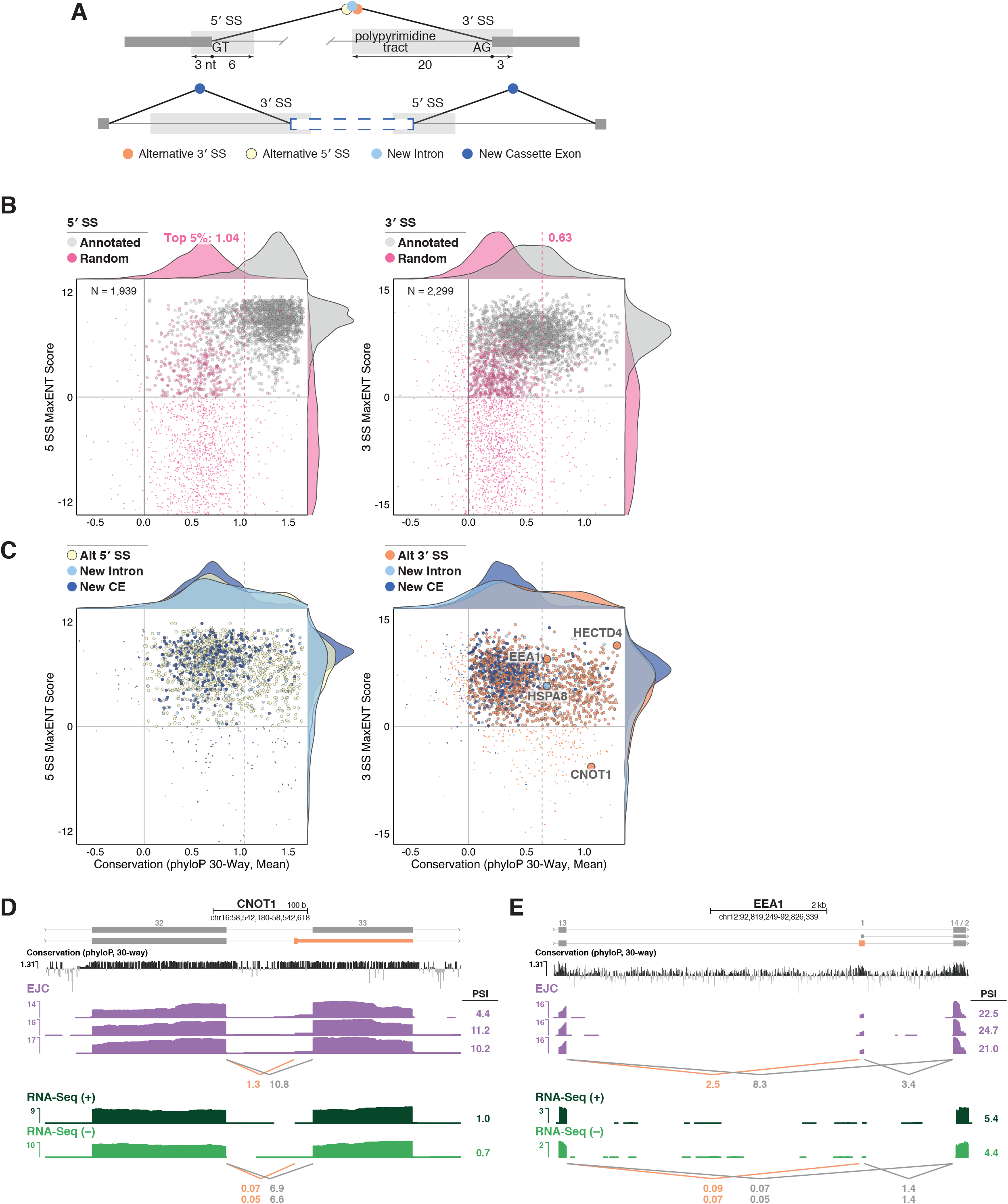
(A) Regions used to calculate MaxEnt and mean conservation scores surrounding unannotated alternative 3′ and 5′ splice sites and new introns (top) or new cassette exons (bottom). (B and C) Scatterplots comparing MaxEnt scores to mean conservation scores (phyloP, 30-way) at 5′ (left) or 3′ (right) splice sites for (B) annotated and random or (C) observed unannotated events. Smaller points are used to represent splice sites with either score lower than 0 as these may result from splicing noise. Annotated splice sites were downsampled by random selection (5′, N = 2,268; 3′, N = 2,693; same as unannotated splice site numbers in C) from the 159,335 observed in our libraries. Supplemental Figure 4A shows the same plot for all observed annotated splice sites. (B) also contains 2,268 random GT-containing (left) and 2,693 random AG-containing (right) sites; identical plots for four additional sets of randomized locations are shown in Supplemental Figure 4B. The top 5% mean conservation scores of random sites is indicated and marked by a dashed line. Genes for which genome-browser tracks are shown in panels D and E and Supplemental Figures 5A and B are indicated. (D-E) Genome browser tracks of library coverage across CNOT1 (D) and EEA1 (E). Annotated transcripts are shown in grey and unannotated alternative 3′ splice site use in orange. Conservation tracks and annotations are as in Figure 1B-D.

### New evolutionarily-conserved poison cassette exons

Having found examples of new AS-NMD isoforms generated by unannotated 3′ splice sites, we were interested to investigate which of the new cassette exons identified here might also function in this capacity. Of our 413 new cassette exons (**Figure 3E**), 383 (93%) occurred in protein-coding genes; the remainder occurred in pseudogenes and ncRNAs. Based on the data in **Figure 1**, poison exons should exhibit higher abundance in EJC than in RNA-Seq libraries. Consistent with this, 318/383 (83%) were solely detectable in the EJC libraries, with the remainder averaging 26- and 24-fold higher abundance in EJC than in RNA-Seq libraries treated with (+) or without (–) harringtonine, respectively (**Figure 6A**). Of the 376 new cassette exons detectable in EJC libraries, 70% were frameshifting (i.e., not a multiple of 3 nts long). Individual inspection of the 25 most abundant non-frameshifting exons revealed that 80% contained an in-frame stop codon. Therefore, as expected, the vast majority of our newly identified cassette exons likely function as poison exons.

**Figure 6.**
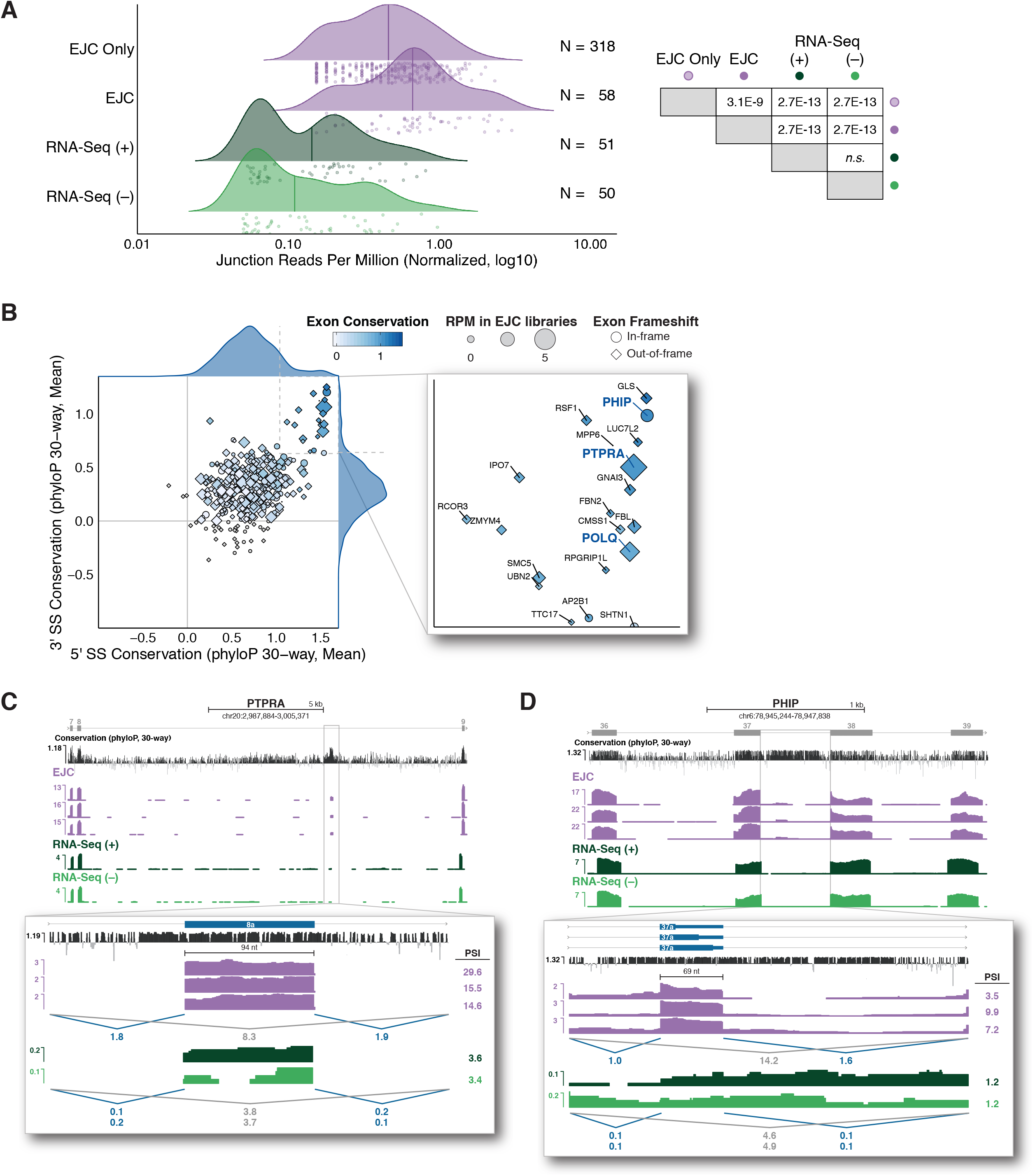
(A) (Left) Density plot comparing junction-spanning read coverage (RPM) for new cassette exons in EJC and RNA-Seq libraries. Line indicates median expression per library and dots represent individual cassette exons. N: number of observed cassette exons per library. (Right) Results of one-way ANOVA and Tukey’s post hoc significance tests comparing the distribution of junction reads in the indicated library types. (B) (Left) Scatterplot comparing mean conservation (phyloP, 30-way) at 5′ and 3′ splice sites of new cassette exons. Exons with scores above 0 at both splice sites are colored (white to dark blue) to indicate mean exon conservation and sized by the number of junction-spanning reads supporting that exon in EJC RIPiT-Seq libraries. Diamonds indicate exons that create a frameshift in the resulting mRNA; circles indicate non-frameshifting exons. (Right) Zoomed view of exons with mean 5′ and 3′ splice site conservation scores above 1.04 and 0.63, respectively. (B and C) Genome browser tracks of library coverage across new poison cassette exons in PHIP (B) and PTPRA (C). New cassette exons are shown in blue and numbered according to their placement in the major isoform observed in all libraries. Conservation tracks and annotations are as in Figure 1B-D. See Methods for PSI formula.

To assess whether any of the new cassette exons constitute conserved regulatory elements, we calculated mean phyloP 30-way conservation scores across the entire exon. Combining these exon conservation scores (white to dark blue in **Figure 6B**) with the previously calculated 5′ and 3′ splice site conservation scores (**Figure 5B**) revealed a set of 20 previously unannotated cassette exons exhibiting both high internal (phyloP score ≥ 1) and high splice site (≥ 1 for both splice sites) conservation (**Figure 6B right; Supplemental Table 3**). Among these, the most highly represented in our datasets was a new 94 nt exon within intron 8 of the 22-intron protein tyrosine phosphatase, receptor type A (PTPRA) gene (**Figure 6C**). Reminiscent of the conserved poison exons in TRA2B and U2AF2 (**Figure 1B, C**), inclusion of (PTPRA) exon 8a was readily observable in the EJC libraries, but nearly undetectable in the RNA-Seq libraries (**Figure 6C**). Other high abundance examples were a 103 nt exon in intron 3 of the 29-intron DNA Polymerase Theta (POLQ) gene (**Supplemental Figure 5C**) and a 69 nt exon in intron 37 of the 39-intron pleckstrin homology domain interacting protein (PHIP) gene (**Figure 6D**). Although PHIP exon 37a does not frameshift, it does contain three highly-conserved in-frame stop codons (**Figure 6D, bottom**). Thus all of the new evolutionarily-conserved cassette exons identified here likely function as poison exons to regulate protein expression from their host gene.

## Discussion

Here we demonstrate that deep sequencing of transcripts in pre-translational RNPs provides a means to identify and quantify mRNA isoforms underrepresented in or absent from RNA-Seq libraries due to their rapid elimination by translation-dependent mRNA decay. We captured this pre-translational population by tandem immunoprecipitation (RIPiT) [12] of two core EJC proteins. EJCs are stably deposited upstream of exon junctions late in the pre-mRNA splicing process, and EJCs in 5′ UTRs and coding regions (∼98% of all) are necessarily removed during the first or “pioneer” round of ribosome transit. Thus the EJC provides an excellent handle by which to enrich for fully-processed, but not-yet-translated mRNAs (**Figure 1A**). Our EJC RIPiT-Seq libraries enabled us to identify thousands of new exon junctions not currently annotated in any of four major reference datasets based on RNA-Seq. Many of these new splicing events generate isoforms subject to NMD, with some being evolutionarily-conserved AS-NMD regulatory events. Thus EJC RIPiT-Seq constitutes a useful method to query the spliced transcriptome without the confounding effects of differential translation-dependent decay of individual mRNA isoforms.

### Measuring flux through AS-NMD pathways

Since its initial description [7, 30, 31], AS-NMD has increasingly emerged as a key post-transcriptional regulatory mechanism [32-34]. Due to their widely different decay rates, however, the flux through the alternative processing pathways resulting in protein-coding and NMD isoforms cannot be captured by traditional RNA-Seq methods. As shown in **Figure 1**, the vast majority of TRA2B and U2AF2 transcripts present in RNA-Seq libraries are protein coding isoforms. The EJC RIPiT-Seq libraries, however, tell a very different story. For both TRA2B and U2AF2, the predominant pre-translational isoform is the poison-exon-included isoform, with poison exon PSIs averaging 88.4% and 57.5%, respectively. Thus, in the cells and growth conditions examined here, alternative splicing flux for both genes strongly favors poison exon inclusion. Similar results were observed for other RNA-binding protein genes known to maintain protein homeostasis by AS-NMD (**Supplemental Figure 1**). Indeed, enrichment of transcripts subject to translation-dependent decay (e.g., isoforms annotated as NMD and NSD) is a general feature of our EJC RIPiT-Seq libraries (**Figure 2**).

One can increase the abundance of transcripts subject to translation-dependent decay in RNA-Seq libraries by globally inhibiting translation prior to cell lysis [64-65]. Indeed, to enrich for pre-translational RNPs in our RIPiT-Seq experiments, we generally expose cells to a translational inhibitor for 60 minutes prior harvest [9, 12-13, 62]. However, the harringtonine (+) and (–) RNA-Seq data in this study clearly show that this treatment had no discernible effect on either protein-coding or AS-NMD isoform abundance in actively dividing HEK293 cells (**Figures 1-2 and Supplemental Figures 1-2**). Another recent study provided whole cell RNA-Seq data for HeLa cells either not subjected to translation inhibition or incubated with 100 mg/ml cycloheximide for 15 minutes or 24 hours [36]. While the 15 minute exposure had almost no effect on protein-coding isoform abundance, the 24 hour exposure did [36] (**Supplemental Figure 6A**). Our analysis of those data revealed that the 15 minute cycloheximide treatment was sufficient to increase AS-NMD isoform abundance, with this increase being even more apparent at 24 hours (**Supplemental Figure 6B**). The differences we observed between HEK293 (no detectable increase in AS-NMD isoform abundance after 60 min harringtonine treatment) and HeLa (clearly detectable increase in AS-NMD isoform abundance after 15 min cycloheximide treatment) could be due to use of different translation inhibitors or to different mRNA synthesis and translation-dependent decay kinetics between the two cell types. Nonetheless, these data illustrate the complexities of trying to assess synthetic flux through AS-NMD pathways using translation inhibition and RNA-Seq alone.

### Identification of previously unannotated conserved splicing events

A major goal for this study was to assess the utility of EJC RIPiT-Seq libraries for identifying sites of exon ligation underrepresented in traditional RNA-Seq libraries. As illustrated in **Figure 3A**, there remain substantial differences in exon ligation events annotated in RefSeq and Ensembl/GENCODE. Further, not all exon ligation events annotated in these reference sets were returned by the CHESS pipeline, the deepest analysis of RNA-Seq to date. Here we identified thousands of exon junctions not currently annotated in RefSeq, Ensembl/GENCODE or CHESS (**Figure 3C**). Whereas the majority of these events occur at sites lacking splice site conservation (**Figures 5 and 6**) and so likely constitute splicing noise, hundreds exhibit high sequence conservation among mammals. Among this conserved set, the majority display features expected to generate an AS-NMD isoform (i.e., frameshift or in-frame stop codon).

### New poison exons regulate genes linked to cancer

It has now been well established that changes to pre-mRNA splicing patterns can drive both cancer initiation and cancer progression [37, 38]. Thus it is of particular note that three of the most conserved, high-abundance AS-NMD events discovered here are poison cassette exons in PTPRA, PHIP, and POLQ (**Figure 6**). All three genes have been linked to poor cancer prognosis when overexpressed [39-44]. While protein overexpression in cancer often results from gene duplication or transcriptional dysregulation, decreased flux through a splicing pathway leading to poison exon inclusion would have the same effect. Previous studies examining the links between NMD and cancer have focused mainly on loss of tumor suppressor genes due to increased NMD [45, 46] or the advantageous effects of NMD in eliminating mRNA isoforms encoding neoepitopes that would otherwise be recognized by the immune system [47]. But our findings suggest that decreased poison exon inclusion should also be considered as a contributor to the mechanisms underlying cancer. An obvious means to alter splicing flux is a cis-acting mutation that disrupts splice site recognition and, thereby, poison exon inclusion. Although our examination of The Cancer Genome Atlas (Release 19) [48] database revealed no instances of splice site mutations associated with any of the new conserved poison cassette exons documented here, this possibility should certainly be considered in future hunts for cancer-promoting mutations. Of note, current “exome” sequencing generally captures only DNA covering and surrounding annotated exons [49]. Therefore, the unannotated cassette exons we identified here are likely absent from most DNA sequencing databases.

## Conclusions

Sequencing of post-splicing, pre-translational mRNPs provides a powerful approach to identify and quantify transient species that undergo rapid translation-dependent decay and are therefore under-represented in or completely absent from standard RNA-Seq libraries. The data here constitute just one snapshot of AS flux in HEK293 cells growing under optimal conditions. Future studies examining EJC RIPiT-Seq libraries from more diverse biological samples will undoubtedly lead to discovery of even more previously undocumented AS-NMD pathways. Examination of how flux through such pathways changes in response to changing cellular conditions will increase our general understanding of how post-transcriptional mechanisms regulate protein abundance.

## Methods

### Deep sequencing libraries

EJC RIPiT-Seq libraries were downloaded from the NCBI GEO GSE115788 (specifically, samples GSM3189985, GSM3189986, and GSM3189987). These libraries were generated from 200-550 nt fragments by 3′ adaptor ligation and reverse transcription. Paired-end sequencing (150 nt reads) on the Illumina NextSeq platform resulted in 18-24 million mate pairs per replicate [13]. For RNA-Seq libraries, HEK293 cells were grown as in [13] and subjected (+) or not (–) to a one hour treatment with harringtonine (2 ng/mL) prior to cell harvest. RNA was isolated in TRIzol using the Direct-zol RNA Kit (Zymo Research, R2062). Deep sequencing libraries were prepared from three biological replicates per condition using the KAPA RNA HyperPrep Kit with RiboErase (HMR) (Roche, 08098131702) with a modified protocol as follows. Isolated RNA (7 ug; Quantified using Qubit RNA Broad Range Assay Kit (Thermo Fischer Scientific, Q10210) was first treated with Turbo DNase using the standard protocol (Thermo Fischer Scientific, AM2239). Treated RNA (1 ug) was then used as input at the rRNA depletion step in the KAPA kit protocol. To generate fragments of similar size to EJC RIPiT-Seq libraries, fragmentation was carried out for 5 minutes at 94°C and samples immediately chilled on ice. Following seven amplification cycles using the dual index adapters (5 uL of a 1.5 uM per uL dilution; final concentration of 10 nM) from the KAPA Dual-Indexed Adapter Kit (KK8722) (Roche, 08278555702), PCR products containing 200 to 550 nt inserts were size selected using SPRIselect beads (Beckman Coulter Life Sciences, B23318) and quantified using the Qubit dsDNA Broad Range Assay Kit (Thermo Fischer Scientific, Q32850). As each library contained a unique barcode, all libraries (3 biological replicates + or - harringtonine treatment) were mixed and sequenced on an Illumina NextSeq 550 instrument using the High Output Kit v2.5 (Illumina, Inc., 20024908). All libraries were loaded at 1.8 pm with 5% PhiX and all data was written to BaseSpace (Illumina, Inc.).

### Library processing and alignment

Read counts for unprocessed libraries and for the individual processing steps detailed below are provided in **Supplemental Table 1**. Prior to alignment, adaptor sequences and long stretches (≥ 20 nt) of adenosines were trimmed from read 3′ ends. All libraries were filtered for reads aligning to repeat regions, as defined by RepeatMasker [15], using STAR v2.5.3a [14]. Remaining reads were aligned with STAR on two-pass mode to the human genome, release 93 [3]. This alignment allowed a maximum of 3 mismatches per pair and highly penalized deletions and insertions. Mapped reads were then filtered for low mapping quality (MAPQ < 5) and/or duplicated reads, identified with the MarkDuplicates tool (Picard v2.17.8) [50].

### RNA isoform quantification

RNA isoform abundances were determined using Kallisto (v0.44.0) [51], using only reads that passed the filtering and alignment steps described above. Transcript biotypes (i.e., “protein-coding”, “nonsense-mediated decay”, etc.) and intron counts used to categorize transcripts throughout **Figure 2, Supplemental Figure 2** and **Supplemental Figure 6A** are based on the transcriptome annotation from Ensembl (GRCh38.p12) [3].

### Junction identification pipeline

The custom bioinformatics pipeline designed for our annotated and unannotated junction analysis (**Figures 3 – 6**) is shown in detail in **Supplemental Figure 3A**. Transcriptome annotation files from RefSeq (hg38) [20], Ensembl (GRCh38.p12) [3], GENCODE (v29) [21], and CHESS (v2.1) [22] were combined to create a comprehensive reference file of all annotated introns (**Supplemental Table 2**). Any junction that appears in our libraries but is not annotated in one of the aforementioned transcriptomes is referred to as “unannotated.”

To identify unannotated exon junctions, all reads with CIGAR strings containing an “N” operation were isolated and compared to the annotated intron reference file using Bedtools intersect [52]. Reads with alignment gaps not matching the length or location of a known intron were considered the result of potential unannotated splicing events. These junctions were further filtered based on the following criteria: (i) overlap with a known gene; (ii) reads must have ≥15 nt aligned on both sides of the potential junction; (iii) present in all replicates of any library type; (iv) GT and AG dinucleotides at the 5′ and 3′ splice sites, respectively; and (v) mean read count ≥ 2 per library type.

### Nearest annotated splice site analysis

For analysis of new splicing events near annotated exons (**Figure 4**), each unannotated 5′ splice site was paired with its nearest annotated 5′ splice site based on the 3′ splice site used in both splicing events. Similarly, each unannotated 3′ splice site was paired with its nearest annotated 3′ splice site based on the 5′ splice site used in both splicing events. The number of available GT and AG dinucleotides at nucleotide positions −30 to +30 surrounding each annotated splice site in this unannotated/annotated paired dataset were counted to determine the frequency of potential splice sites in the relevant region.

### Splice site strength and conservation

Splice site strength and mean conservation scores for annotated and unannotated splice sites were calculated using MaxEntScan [29] and phyloP 30-way basewise conservation scores [28] (**Figure 5A**). Random sequences of the appropriate length (9 nts for 5′ splice sites and 23 nts for 3′ splice sites) and internal to annotated genes were obtained from the hg38 annotation file [3] using the Bedtools random function [52]. Only those random sequences containing a GT at positions 4 and 5 or an AG at positions 19 and 20 were used to calculate MaxENT and conservation scores for comparison to 5′ and 3′ splice sites, respectively.

### Plotting and data visualization

Data visualization was performed in R [53] using ggplot2 [54], ggrepel [55], UpSetR [56], ggseqlogo [57], eulerr [58], and ggridges [59] software packages. The UCSC Genome Browser [60, 61] was used to view library tracks and to create transcript figures throughout the manuscript.

### Splice site usage calculations

The following equations were used to calculate RPM, PSI, PSO, and PIR values throughout the manuscript. NMD and PC refer to Nonsense-Mediated Decay and Protein Coding isoforms, respectively.

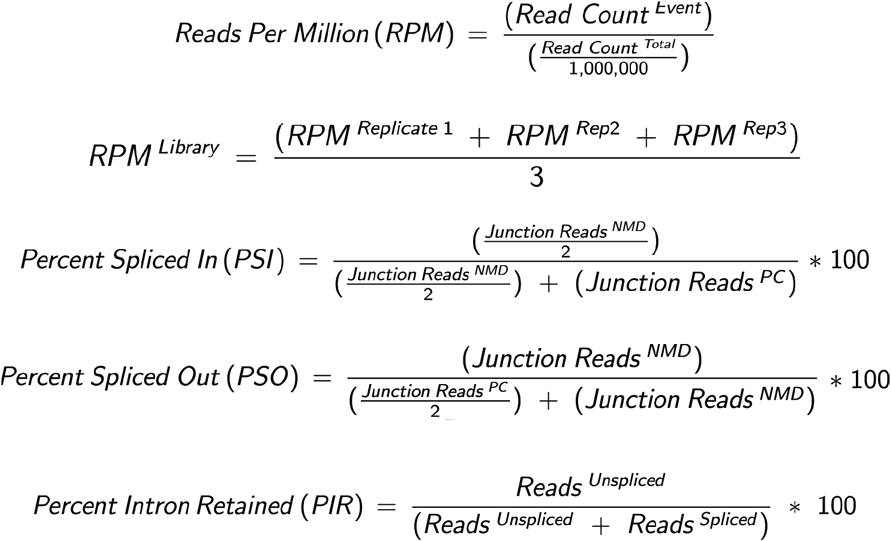

## List of abbreviations

AS: alternative splicing
RNA-Seq: deep sequencing of RNA
NMD: nonsense-mediated decay
NSD: non-stop decay
AS-NMD: alternative splicing linked to NMD
EJC: exon junction complex
cEJCs: canonical exon junction complex
nEJCs: noncanonical exon junction complex
RIPiT-Seq: RNA:protein immunoprecipitation in tandem followed by deep sequencing
PSI: percent spliced in
PSO: percent spliced out
PIR: percent intron retained
CHESS: Comprehensive Human Expressed SequenceS
RPM: reads per million
TPM: transcripts per million

## Declarations

### Additional files

#### Additional file 1

Includes Figures S1-S5 and Table S1. **Figure S1**. Comparison of biological replicates; additional examples of AS-NMD transcript coverage. **Figure S2**. Comparison of EJC and RNA-Seq libraries coverage across additional transcript biotypes. **Figure S3**.

Representation of bioinformatics pipeline to classify annotated and unannotated exon junctions. **Figure S4**. Comparison of MaxENT and conservation scores for annotated and randomly located splice sites; additional examples of conserved unannotated splicing events. **Figure S5**. Additional example of an unannotated AS-NMD cassette exon. **Figure S6**. Analysis of previously published RNA-Seq libraries made from HeLa cells treated with cycloheximide for 0 minutes, 15 minutes, or 24 hours [36]. **Table S1**. Sequencing and alignment information for each replicate of the analyzed libraries.

#### Additional file 2

Includes Table S2. **Table S2**. Comprehensive file of all intron locations annotated in RefSeq (hg38), Ensembl (GRCh38.p12), GENCODE (v29), and CHESS (v2.1).

#### Additional file 3

Includes Table S3. **Table S3**. Locations and characteristics of highly conserved unannotated exon junctions identified in this study.

## Acknowledgments

We thank Athma Pai, Harleen Saini and Guramrit Singh for critical reading of the manuscript. We thank Weijun Chen for technical advice. We thank Wei Zheng and Chiao-wen Hsiao for advice on statistical analysis.

## Availability of data and materials

The RIPiT-Seq datasets analyzed in this study were downloaded from NCBI GEO under accession number GSE115788 (specifically, samples GSM3189985, GSM3189986, and GSM3189987). The accession number for the RNA-Seq raw fastq files reported in this paper is GSE150228.

## Authors’ contributions

M.J.M conceived of the project. C.K. and M.J.M were responsible for all data analysis and manuscript writing. S.D. and A.A.B. constructed the RNA-Seq libraries and participated in manuscript editing. M.M. provided technical advice regarding the EJC RIPiT-Seq libraries and participated in manuscript editing.

## Ethics approval and consent to participate

Not applicable.

## Consent for publication

Not applicable.

## Author details

M.J.M. is a member of the National Academy of Science (USA) and a fellow of the American Association of Arts and Sciences.

## Figure Legends

*Supplemental Figure 1*

(A) Scatterplots comparing transcripts per million (TPM) between replicates of individual library types. R: Pearson’s correlation. (B-D) Genome browser tracks of library coverage across individual genes (grey: protein-coding isoform(s); blue: NMD isoforms) that utilize a poison cassette exon (B, SRSF6), exon skipping (C, SNRPA1) or 3′ UTR introns (D, PRPF4B) to generate NMD isoforms. Shown are all three EJC RIPiT-Seq replicates and replicate 1 for both (+) and (–) harringtonine RNA-Seq libraries. Conservation tracks show phyloP basewise scores derived from Multiz alignment of 30 vertebrate species. Numbers below tracks indicate mean reads per million (RPM) spanning each exon junction. Numbers on the right in B and C are percent spliced in (PSI) and percent spliced out (PSO) values, respectively; PSI and PSO values for RNA-Seq libraries are replicate means. See Methods for PSI and PSO formulae.

*Supplemental Table 1*

Sequencing and alignment information for each replicate of all analyzed libraries.

*Supplemental Figure 2*

(A) Scatterplots comparing TPMs between EJC RIPiT-Seq and RNA-Seq (+) libraries for non-stop decay (left), retained intron (middle), and processed transcript (right). N: Number of detected transcripts (out of all annotated transcripts of that type). Dashed black line: x=y. (Inset) Distribution of spliced and unspliced transcript expression of each transcript biotype. (B) Ratios of transcript abundance between two indicated library types, binned based on transcript biotype [3]. For A and B: Results of one-way ANOVA and Tukey’s post hoc significance tests comparing EJC RIPiT-Seq to RNA-Seq libraries are indicated as *P<0.05, **P<0.01, ***P<0.005, ****P<0.0001.

*Supplemental Figure 3*

(A) Schematic of library processing steps used to identify and analyze reads at annotated and unannotated junctions. Full details of each step are in Results and Methods.

*Supplemental Table 2*

List of all introns previously annotated by RefSeq (hg38) [20], Ensembl (GRCh38.p12) [3], GENCODE (v29) [21], and/or CHESS (v2.1) [22]. Table includes information on intron location, length, strand, transcript ID (if available, [3]), annotation origin, and normalized (and raw) counts of junction reads per replicate of each library type.

*Supplemental Figure 4*

(A) Scatterplots comparing the MaxEnt score to mean conservation score (phyloP, 30-way) at 5′ (left) or 3′ (right) splice sites for all annotated junctions (N = 159,335). (B) Scatterplots comparing the MaxEnt score to conservation (phyloP, 30-way) at 5′ (left) or 3′ (right) splice sites for multiple sets (N = 5) of randomly selected sequences (5′, N = 2,268; 3′, N = 2,693). (C) Table of calculated mean differences and standard errors (SEM) between unannotated splice sites and annotated or random sequence MaxENT and phyloP 30-way conservation scores; 95% confidence intervals (CI) and p-values are from independent (unpaired) t-tests.

*Supplemental Table 3*

List of highly conserved unannotated splicing events (see **Results** for conservation score cut-offs). Table includes information on exon junction locations (coordinates and transcript features), MaxENT and conservation scores, calculated PIR values, and normalized (and raw) counts of junction reads per replicate of each library type.

*Supplemental Figure 5*

(A-B) Genome browser tracks of library coverage across HSPA8 (A) and HECTD4 (B). Annotated transcripts are shown in grey, unannotated alternative 3′ splicing events in orange, and unannotated introns in light blue. Conservation tracks represent phyloP basewise scores derived from Multiz alignment of 30 vertebrate species, as well as 100 vertebrate species in (B). Numbers below tracks indicate mean reads per million (RPM) spanning each exon junction. Numbers to the right in A and B are percent intron retention (PIR) and percent spliced in (PSI) values, respectively; PSI and PIR values for RNA-Seq libraries are replicate means. See Methods for PSI and PIR formulae. The translated protein sequences of both the annotated and unannotated transcripts are provided in (B). (C) Genome browser tracks of library coverage across the new cassette exon in POLQ. The new cassette exon is shown in blue and numbered according to its placement in the major isoform observed in all libraries. Conservation tracks and annotations are as in Figure 1B-D. See Methods for PSI formula.

*Supplemental Figure 6*

(A) Volcano plots comparing cumulative protein-coding isoform abundance differences between untreated HeLa cells and HeLa cells treated with cycloheximide for 15 minutes (left) or 24 hours (right) using deep sequencing libraries from [36]. Each dot is a gene. Dashed lines indicate cut-offs at log_2_FoldChange > |2| and log_10_padj > 0.005. Labeled transcripts: The sole gene meeting cutoff criteria at 15 min (BAG5) plus those genes highlighted in [36] Supplemental Figure 7 found to be up-regulated (blue) or down-regulated (red) in our analysis. (B) Genome browser tracks of library coverage across the individual genes (grey: protein-coding isoform(s); blue: NMD isoforms) highlighted in Figure 1B-D and Supplemental Figure 1B-D. Shown are replicate 1 for RNA-Seq libraries generated from HeLa cells either not subjected to translation inhibition or incubated with 100 mg/ml cycloheximide for 15 minutes or 24 hours [36]. Numbers to the right are percent spliced in (PSI) and percent spliced out (PSO) values, respectively; PSI and PSO values for RNA-Seq libraries are replicate means. See Methods for PSI and PSO formulae.

