## Supplemental Figures for "Deep sequencing of pre-translational mRNPs reveals hidden flux through evolutionarily conserved AS-NMD pathways"

### Supplemental Figure 1

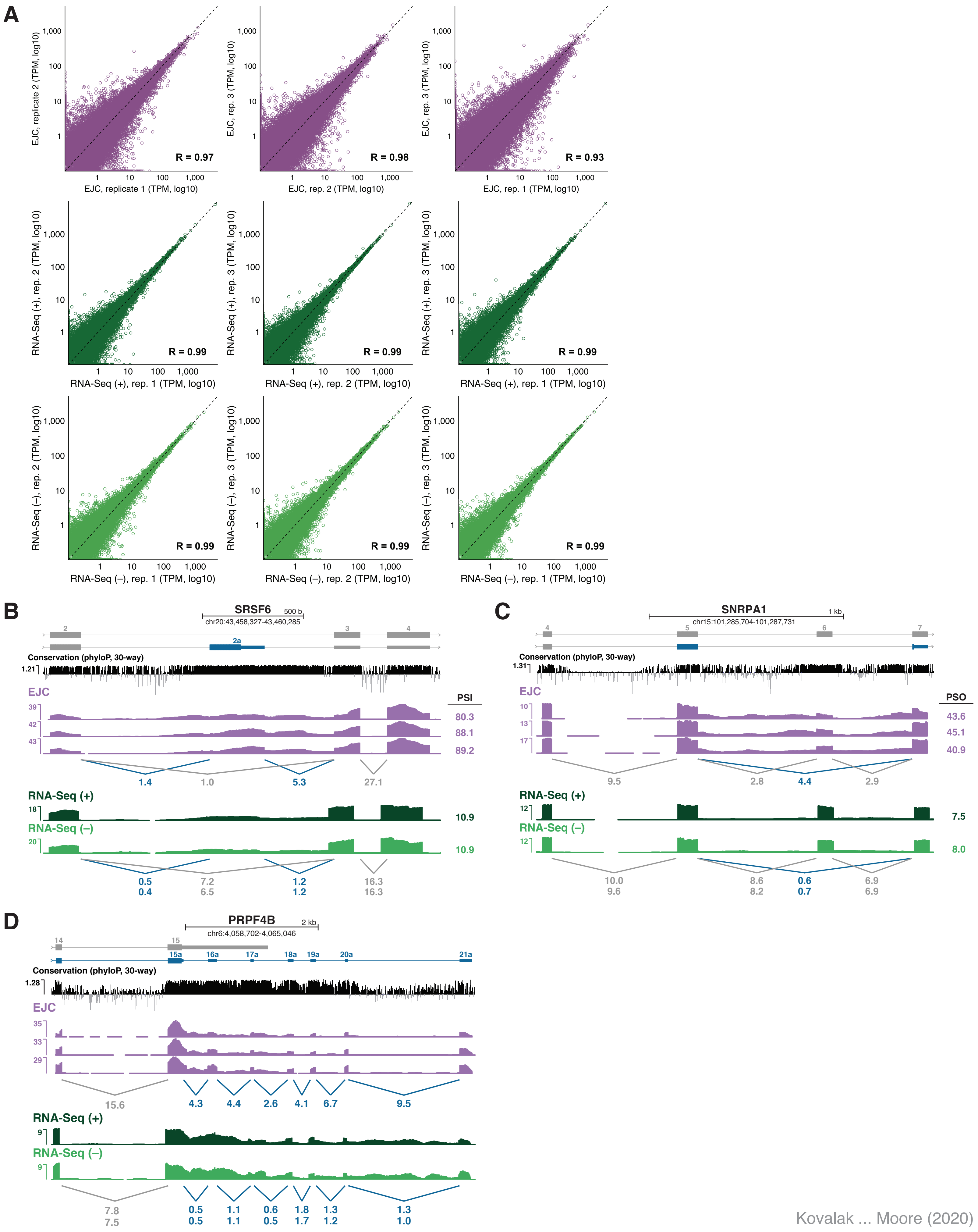


Kovalak ... Moore (2020)

### Supplemental Figure 2

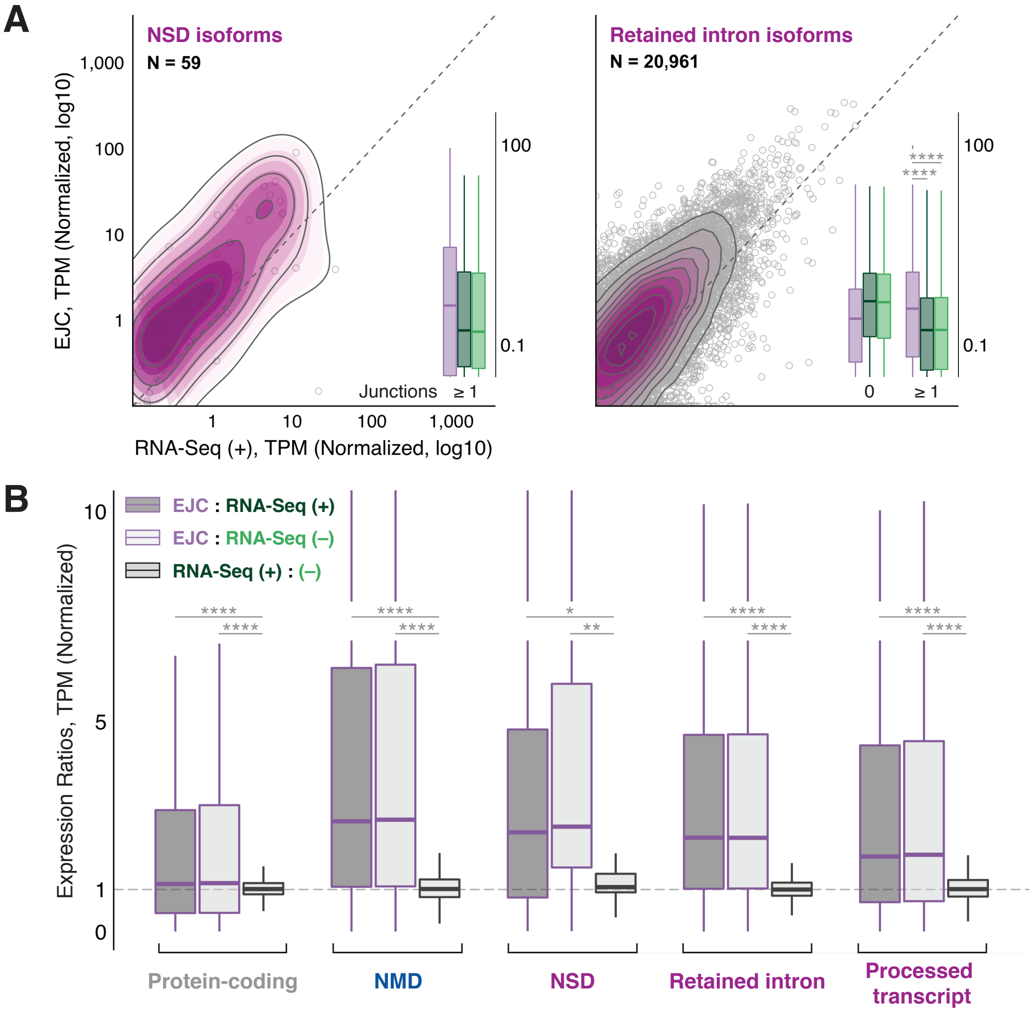

#### Supplemental Figure 3

A

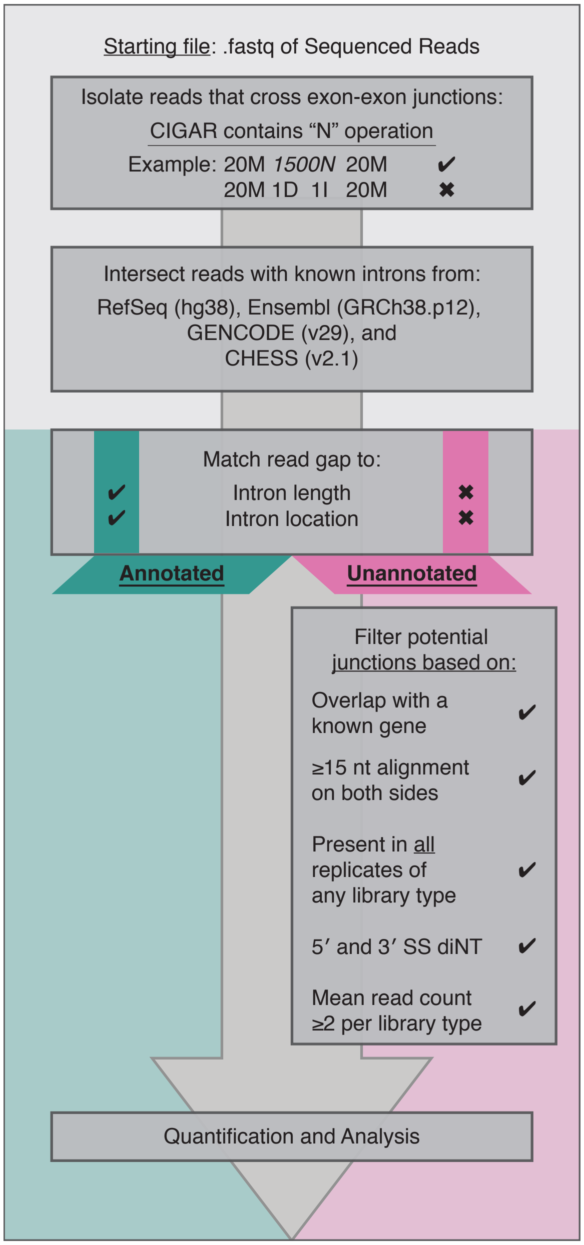

### Supplemental Figure 4

A

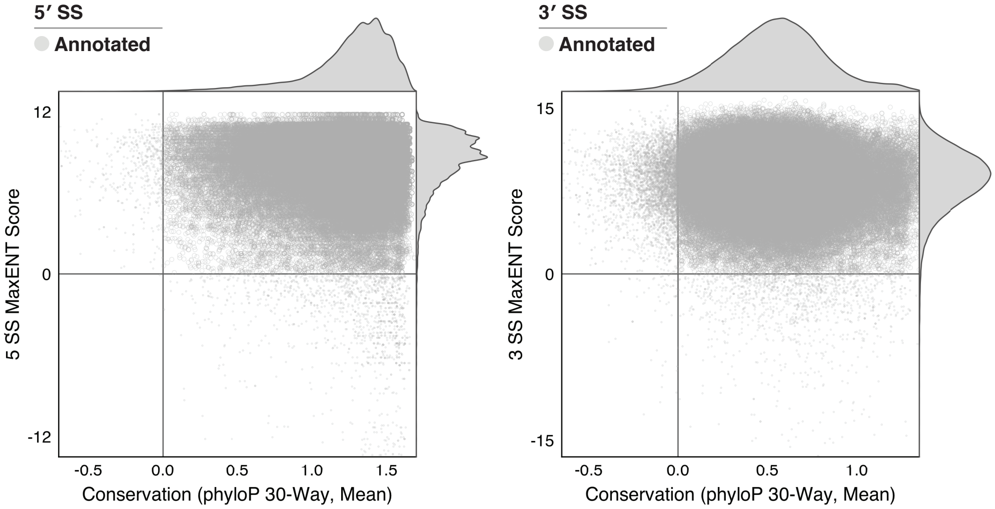


B

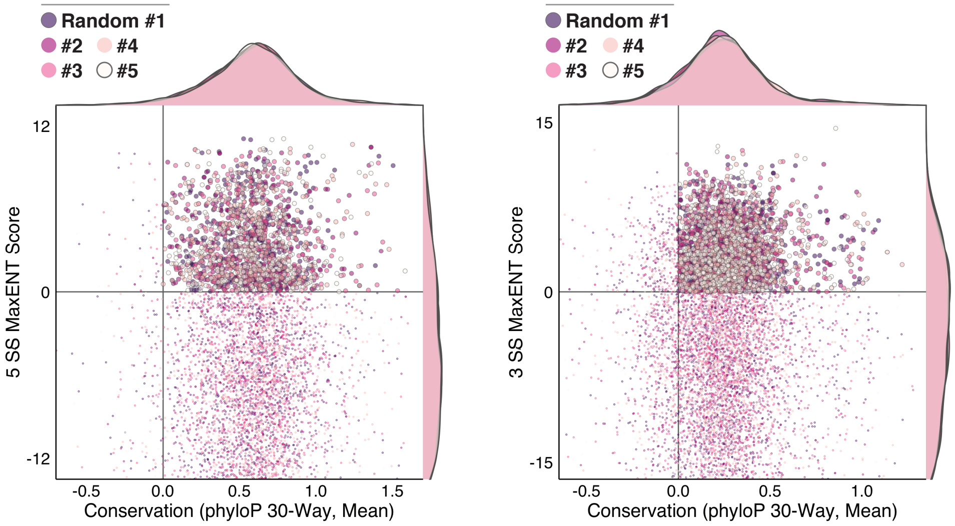


C

| | | To Annotated | | | To Random | | |
|---|---|---|---|---|---|---|---|
| **5′ SS** | | Mean diff. ± SEM | 95% CI | p-value | Mean diff. ± SEM | 95% CI | p-value |
| **MaxENT** | Alt 5′ SS | 2.31 ± 0.11 | [2.05, 2.56] | < 2.2e-16 | -14.60 ± 0.29 | [-15.07, -14.13] | < 2.2e-16 |
| | New Intron | 3.62 ± 0.21 | [2.78, 4.60] | 2.7e-08 | -13.29 ± 0.76 | [-14.22, -12.36] | 2.6e-08 |
| | New CE | 0.94 ± 0.10 | [0.71, 1.17] | 1.7e-03 | -15.97 ± 0.35 | [-16.42, -15.52] | 4.2e-08 |
| **Cons.** | | 0.44 ± 0.01 | [0.41, 0.47] | < 2.2e-16 | -0.25 ± 0.01 | [-0.28, -0.23] | < 2.2e-16 |
| | | 0.47 ± 0.02 | [0.40, 0.54] | 2.6e-08 | -0.22 ± 0.03 | [-0.30, -0.15] | 2.6e-08 |
| | | 0.55 ± 0.01 | [0.52, 0.58] | 4.2e-08 | -0.14 ± 0.01 | [-0.17, -0.12] | 4.2e-08 |
| **3′ SS** | | | | | | | |
| **MaxENT** | Alt 3′ SS | 3.66 ± 0.12 | [3.40, 3.92] | < 2.2e-16 | -11.32 ± 0.27 | [-11.77, -10.86] | < 2.2e-16 |
| | | 4.38 ± 0.27 | [3.22, 5.53] | 3.2e-08 | -10.60 ± 0.80 | [-11.81, -9.39] | 3.2e-08 |
| | | 1.21 ± 0.12 | [1.14, 1.61] | 8.2e-08 | -13.78 ± 0.37 | [-13.66, -12.86] | < 2.2e-16 |
| **Cons.** | | 0.01 ± 0.01 | [-0.01, 0.03] | 0.31 | -0.29 ± 0.01 | [-0.31, -0.27] | < 2.2e-16 |
| | | 0.17 ± 0.02 | [0.11, 0.23] | 3.2e-08 | -0.13 ± 0.02 | [-0.19, -0.07] | 5.8e-08 |
| | | 0.22 ± 0.01 | [0.20, 0.24] | < 2.2e-16 | -0.08 ± 0.01 | [-0.10, -0.06] | 1.8e-11 |

### Supplemental Figure 5

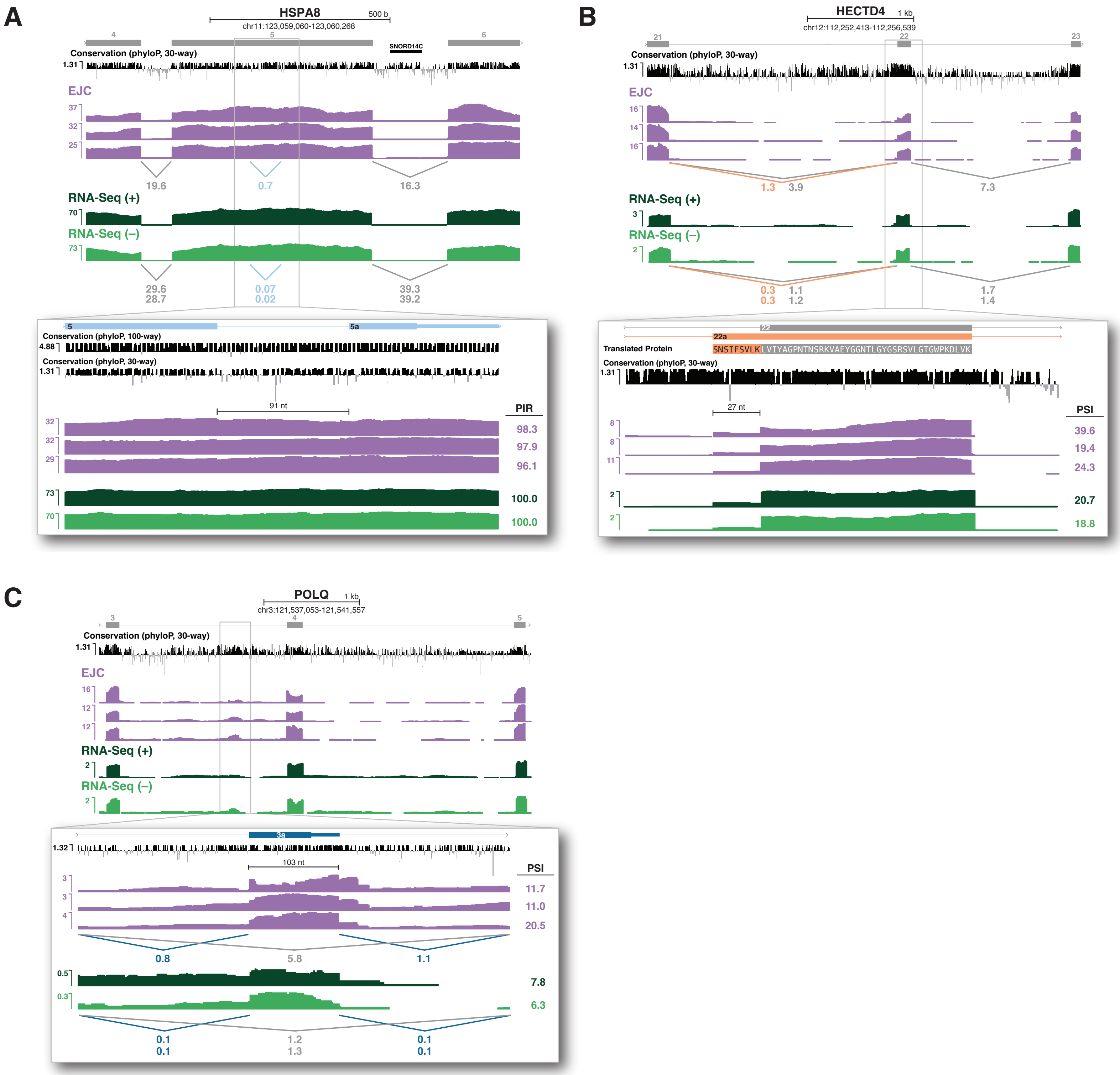

### Supplemental Figure 6

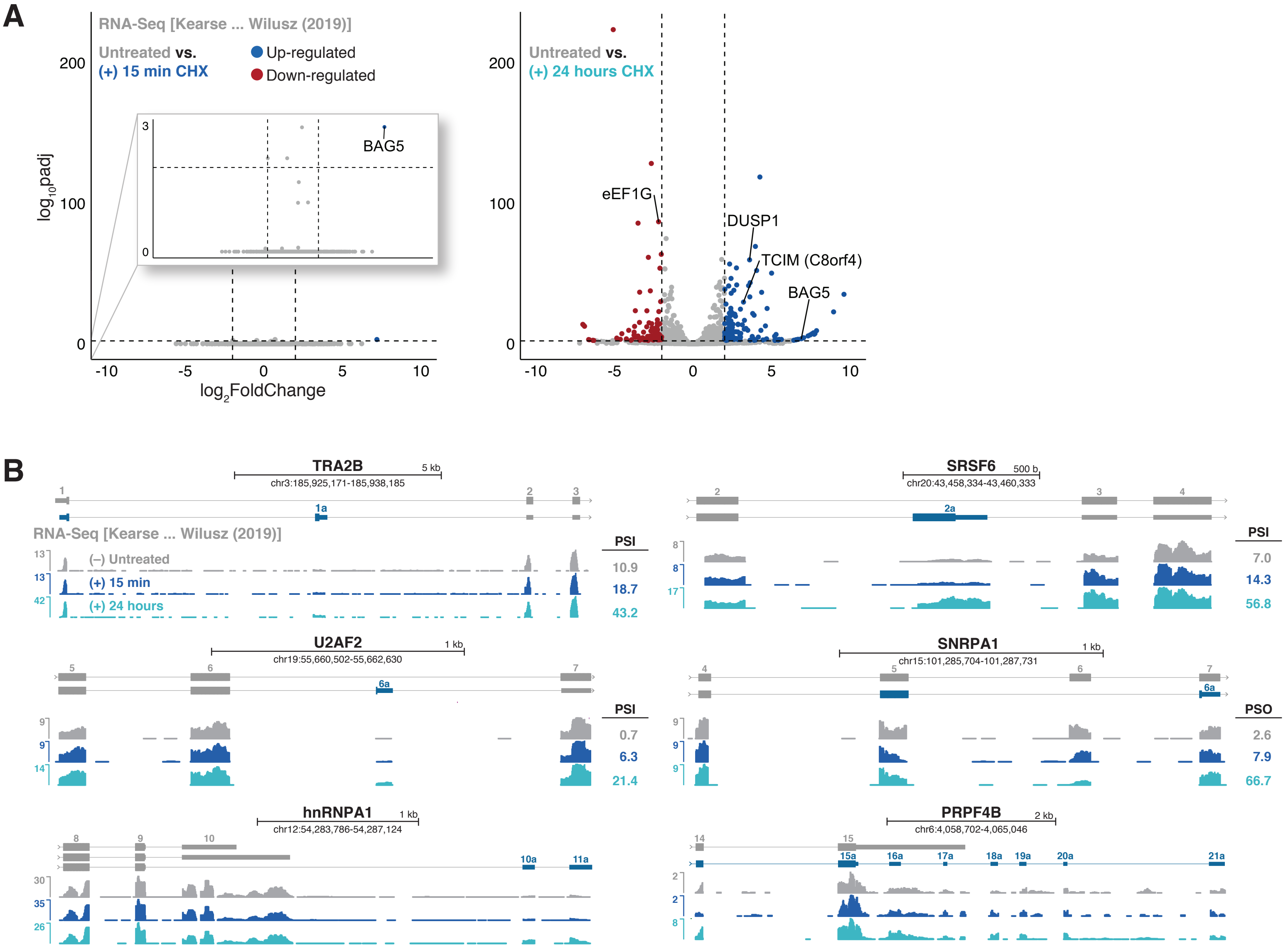

### Supplemental Table 1

| Library | | | Sequencing | | | Alignment | | | | Analysis |
|---|---|---|---|---|---|---|---|---|---|---|
| *Name* | *Harringtonine* | *Replicate* | *Type* | *Insert Size* | *Sequenced Pairs* | *Repeats* | *Aligned Pairs* | *MAPQ ≥ 5* | *Unique Pairs* | *Spliced Reads* |
| **EJC** | + | 1 | Paired End, 150bp | 200-550 | 19 Million | – 3 M | 6 M | 5.6 M | 5.1 M | 3.3 M |
| | | 2 | | | 25 M | – 4 M | 10 M | 9.0 M | 8.2 M | 5.6 M |
| | | 3 | | | 23 M | – 4 M | 7 M | 6.6 M | 6.1 M | 4.3 M |
| **RNASeq** | + | 1 | Paired End, 150bp | 200-550 | 92 M | – 16 M | 67 M | 63.2 M | 56.2 M | 35.8 M |
| | | 2 | | | 84 M | – 15 M | 60 M | 56.6 M | 50.2 M | 34.9 M |
| | | 3 | | | 87 M | – 16 M | 62 M | 58.0 M | 51.2 M | 35.5 M |
| **RNASeq** | – | 1 | Paired End, 150bp | 200-550 | 91 M | – 17 M | 76 M | 61.6 M | 54.5 M | 36.8 M |
| | | 2 | | | 92 M | – 17 M | 64 M | 59.9 M | 53.2 M | 33.9 M |
| | | 3 | | | 93 M | – 17 M | 67 M | 62.7 M | 55.6 M | 34.7 M |

### Supplemental Table 2

See file: Supp_Table_2.txt

### Supplemental Table 3

See file: Supp_Table_3.txt
